# Isoliquiritigenin attenuated cognitive impairment, cerebral tau phosphorylation and oxidative stress in a streptozotocin-induced mouse model of Alzheimer’s disease

**DOI:** 10.1101/2024.11.28.625892

**Authors:** Zhi Tang, Ting Sha, Yuanshang Wang, Yan Xiao, Yuanting Ding, Ruiqing Ni, Xiaolan Qi

**Affiliations:** Key Laboratory of Endemic and Ethnic Diseases, Ministry of Education & Key Laboratory of Medical Molecular Biology of Guizhou Province, Guizhou Medical University, Guiyang, China; The Affiliated Baiyun Hospital of Guizhou Medical University, Guiyang, China; Institute for Regenerative Medicine, University of Zurich, Zurich, Switzerland

**Keywords:** Alzheimer’s disease, ERK, GSK-3β, isoliquiritigenin, mitochondria, mTOR, oxidative stress, streptozotocin, tau

## Abstract

**Introduction:** Tau hyperphosphorylation, mitochondrial dysfunction and oxidative stress play important roles in Alzheimer’s disease (AD). Isoliquiritigenin, a natural flavonoid isolated from the root of liquorice, has been shown to exert inhibitory effects on oxidative stress. Here, we assessed the neuroprotective effects of isoliquiritigenin on a streptozotocin-injected mouse model.

**Method:** Molecular docking analysis performed for isoliquiritigenin with mTOR and ERK2. The mice (n = 27, male) were intracerebroventricularly injected with streptozotocin, treated with isoliquiritigenin (intraperitoneal, 2 days) and assessed using the Morris water maze. Oxidative stress, tau phosphorylation, mitochondrial dysfunction and synaptic impairment were evaluated in the cortex and hippocampal tissues of the mice by using biochemical assays and immunofluorescence staining.

**Results:** Isoliquiritigenin treatment mitigated the spatial memory capacity of streptozotocin-injected mice and alleviated tau phosphorylation at Ser396; the production of reactive oxygen species; the mitochondrial fission proteins Mfn1 and Mfn2; neuronal loss; and synaptic impairment (PSD95, SNAP25). Isoliquiritigenin treatment reduced the levels of mTOR Ser2448 and ERK1/2 T202/Y204 and upregulated the level of GSK-3βSer9 in the cortex and hippocampus of streptozotocin-injected mice.

**Conclusion:** In conclusion, our findings suggest that isoliquiritigenin ameliorates streptozotocin-induced cognitive impairment, hyperphosphorylated tau, oxidative stress, mitochondrial dysfunction and synaptic impairment by decreasing mTOR and ERK activity and increasing GSK-3β activity.

## Introduction

Alzheimer’s disease (AD) is the most dominant form of dementia and is pathologically characterized by the accumulation of hyperphosphorylated tau proteins and amyloid-beta (Aβ) plaques and neurodegeneration ^1^. Oxidative stress is characterized by an imbalance between free radical production of reactive oxygen species (ROS) and antioxidant defense and plays a key role in AD ^2^. Elevated levels of ROS are detected in postmortem brain tissues from AD patients and animal models of AD. Hyperphosphorylated tau causes oxidative stress and neurotoxicity and damages microtubule function. In turn, oxidative stress increases tau hyperphosphorylation, forming a vicious cycle. Emerging evidence suggests that aberrant accumulation of activated extracellular signal-regulated kinase (ERK) 1/2, mammalian target of rapamycin (mTOR) and glycogen synthase kinase-3 beta (GSK-3β) signaling pathways in the AD brain is associated with oxidative stress. In addition, mitochondrial defects, impaired brain glucose/energy metabolism and free radical-induced neurodegeneration occur early in AD development. Functional studies have shown disturbances in cerebral glucose metabolism and synaptic density in the medial temporal lobe and frontal cortex of AD patients ^3^, as well as in animal models of AD ^4^. Restoring hippocampal glucose metabolism has recently been shown to rescue cognition across AD pathologies in an animal model ^5^.

Different animal models have been used to study the disease mechanism and treatment for AD. Given the close link between diabetes and AD, the streptozotocin (STZ)-induced rodent model is widely used to induce an insulin-resistant brain state that mimics many aspects of AD pathology ^6^. STZ, a glucosamine-nitrosourea compound, is administered intracerebroventricularly (ICV) and causes insulin signaling malfunction, spatial memory impairment, neuroinflammation, oxidative stress, tau protein phosphorylation, cerebral glucose/energy reduction, and synaptic degeneration ^7–9^. This model is particularly valuable for studying the link between insulin resistance and AD pathogenesis. Compared with transgenic models that focus on familial AD, the STZ model is especially useful for studying ROS and metabolism deficits.

Isoliquiritigenin (ISL) is a natural flavonoid extracted from liquorice (Glycyrrhiza uralensis) ^10^, a medicinal herb ^11, 12^. Among these bioactive flavonoids, ISL is able to penetrate the blood_brain barrier; has various biological effects, including antioxidant, neuroprotective, anti-inflammatory, and antidiabetic activities; and improves mitochondrial function ^13, 14^. In a lipopolysaccharide-induced cognitive impairment rat model, ISL also prevented neuronal damage and cognitive impairment by maintaining antioxidant ability ^15^. In a cerebral ischemia_reperfusion injury model, ISL reduced oxidative stress and ameliorated mitochondrial dysfunction by activating the Nrf2 pathway ^16, 17^. In rats with kainic acid-induced seizures, ISL protects against cognitive impairment by suppressing synaptic dysfunction, neuronal injury, and neuroinflammation ^18^. A previous study reported that ISL alleviated subarachnoid hemorrhage-induced early brain injury in a rat model via the inhibition of neuronal apoptosis, oxidative stress, and brain edema ^19^. ISL effectively reduces rotational behavior and neuroinflammation in a mouse model of Parkinson’s disease ^20^. Previous studies of ISL in AD models have focused on amyloid-beta-induced alterations, such as anti-amyloid neurotoxicity, aggregation, neuroinflammation, NLRP3 inflammasome activation, and ROS production ^11, 21, 22^. The effects of ISL on tau phosphorylation-related alterations and mTOR are not yet clear.

In the present study, we hypothesized that ISL may attenuate STZ-induced detrimental effects on oxidative stress; tau hyperphosphorylation; and mitochondrial, synaptic and cognitive impairment. We evaluated the effects of ISL treatment (intraperitoneal) in an ICV STZ-injected mouse model and the involvement of tau phosphorylation and the GSK-3β, ERK, and mTOR pathways.

## 2 Materials and methods

### 2.1. Network pharmacology

Target prediction and drug-intersecting gene analysis for ISL were conducted via SwissTargetPrediction (http://www.swisstargetprediction.ch/), SuperPred (https://prediction.charite.de/), the BATMAN-TCM database (http://bionet.ncpsb.org.cn/batman-tcm/). Subsequently, keyword searches for ‘Alzheimer’s disease’ were performed in the GeneCards (https://www.genecards.org/) and OMIM (https://www.omim.org/) databases to obtain disease-related targets, excluding duplicate genes. The identified targets of the drug components were mapped against the disease targets, and a Venn diagram was created to obtain intersecting genes. A protein_protein interaction (PPI) network was constructed with the drug_disease intersecting genes using String database (https://string-db.org/). Functional annotation of genes was performed for biological processes (BP), cellular components (CC), and molecular functions (MF) to elucidate the gene functional roles of ISL in the treatment of AD using the drug_disease intersecting genes using DAVID 6.8 and DAVID database (https://david.ncifcrf.gov/summary.jsp)^23^.

### 2.2. Molecular Docking

The 3D structure of ISL was downloaded in SDF format from PubChem, and imported into ChemBio3D Ultra 20.0 for energy minimization, with the minimum RMS gradient set to 0.001. The optimized ISL structure were saved in mol2 format and imported into AutoDockTools-1.5.7 for hydrogenation, charge calculation, charge assignment, and setting of rotatable bonds (The output was saved in “pdbqt” format). The protein structure of MAPK1 (ERK2) (PDB ID: 2Y9Q) was downloaded from the PDB database, and the protein structure of mTOR (AF-P42345-F1) was downloaded from the AlphaFold database. Pymol 3.0.3 was used to remove protein crystal water and the original ligands, after which the protein structures were imported into AutoDockTools (v1.5.7) for hydrogenation, charge calculation, charge assignment, and specification of atom types. The protein binding sites were predicted via the protein plus server, and docking was conducted via AutoDock Vina 1.2.3. PyMOL software (version 3.0.3) and Maestro software (version 13.9.135) were used for analysis and illustration.

### 2.3 Chemicals and antibodies

STZ, trisaminomethane (Tris), radioimmunoprecipitation assay (RIPA), sodium dodecyl sulfate (SDS) buffer and protease inhibitor cocktail were purchased from Sigma Aldrich (USA). For the primary antibodies used in the present study, please refer to **Supplementary Table 1**.

### 2.4 Animals

Twenty-seven C57BL/6J male mice (20–35 g, 2 months of age) were purchased from Sibeifu Biotechnology Co., Ltd. (Beijing, China). Housing and experiments were performed at the Guizhou Experimental Animal Center in China. All the mice were housed in ventilated cages in a climate-controlled room (temperature: 22 ± 2 °C, humidity: 50 ± 5%, 12 h light–dark cycle with lights on at 08:00). Food (safe, sterilized) and water (softened, sterilized) were provided ad libitum. Poplar wood shavings were placed in cages for environmental enrichment. All experimental protocols were approved by the Guiyang Regional Animal Care Center and Ethics Committee (No. 2304628). All the experiments followed ARRIVE guidelines 2.0.

### 2.5 Surgery and treatment

The mice were randomly divided into the control group, STZ group, and STZ + ISL group, with 9 mice per group. In addition, one mouse per group was used for validation of ICV injection and was not used in the following treatment and behavior studies. After the mice were anesthetized with isoflurane and fixed on a stereotactic apparatus, their eyes were coated with petroleum jelly. The heads of the mice were shaved, the bregma was exposed, and the anterior fontanelle was used as the origin, with an injection point medial/lateral (ML): 2.5 mm, anterior/posterior (AP): -1.5 mm. A hole was drilled through the skull, and the needle was inserted 2.5 mm into the right hemisphere hippocampal region (dorsal/ventral (DV): 2.5 mm) (**Fig. 1A**) ^24^. STZ was slowly injected into the right hemisphere of the hippocampus of each mouse at a dose of 3 μL (purity ≥ 95.0%, dissolved in 0.9% sterile saline at a dose of 3 mg/kg). The control group was injected with an equal dose of saline. During the injection, the needle was pushed every 2 minutes for a total of 10 minutes, and after remaining for 2 minutes, it was slowly withdrawn to ensure that the amount of STZ in the local tissue was sufficient. The incision was sutured, and penicillin was applied to the incision to prevent wound infection.

**Fig 1.**
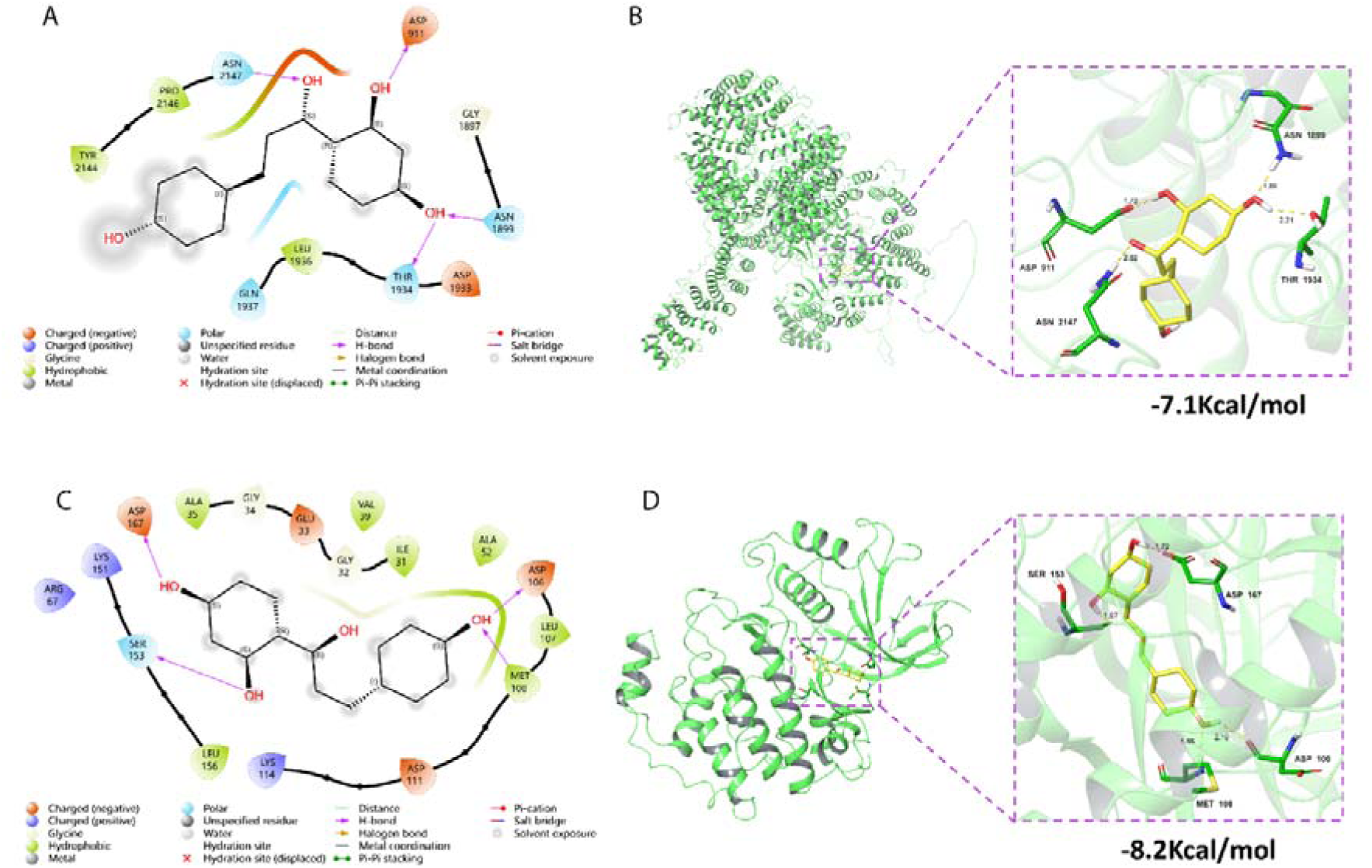
Molecular docking of ISL with ERK2 and mTOR. (**A, B**) 2D and 3D images of the molecular docking of ISL and ERK2. (**C, D**) 2D and 3D images of the molecular docking of ISL and mTOR.

To validate the ICV injection in the brains of the mice, Trypan blue staining was performed on one mouse per group. After ICV injection, the three mice were euthanized, their brains were removed, and 1 mm thick brain slices were obtained with a brain matrix. Slices were incubated in Trypan blue (Beyotime, ST2780-5 g). Photographs of the brain sections of the mice were taken.

The body temperature of the animals was maintained at 25°C throughout the surgery. The mice were allowed to rest for 48 h before ISL treatment. ISL was dissolved in cell-grade dimethyl sulfoxide (DMSO, 10% v/v) and then diluted with polyethylene glycol 300 (PEG300, 40%), Tween-80 (5%), and saline (45%). ISL at a dose of 20 mg/kg was administered intraperitoneally (i.p.) at 48 h, 60 h, 72 h, and 96 h post-surgery, while the control group and the STZ group were given an equal volume of saline (i.p.), followed by subsequent experiments.

### 2.6 Behavioral testing

The mice were allowed to rest for another 2 days after the ISL/saline treatment. The Morris water maze was used to assess the spatial learning ability of the mouse hippocampus ^23, 25^. The circular pool (160 cm diameter and 50 cm height) was filled to a depth of 30 cm with water (25 ± 1 °C) in this study (**Fig. 1B**). The visual cues were positioned above the water level, and the extra maze cues were blocked with a dark curtain. During Morris water maze training, all the mice were subjected to 4 training trials daily for six consecutive days. In each trial, the mice were trained to find a hidden platform (20 cm diameter) submerged 1 cm under the water surface for 60 s. Afterwards, the mice were kept on the platform for 20 s. If these mice could not find the platform within 60 s, they were guided to the platform within 60 s and kept on the platform for 20 s afterwards. On day 5, a spatial probe trial was executed, where the platform was removed. The escape latency, time spent in the target quadrant, number of platform crossings, time to first reach the platform, and distance traveled in the target quadrant of each group of mice were recorded (ANY-maze, USA).

After the behavioral tests, all the mice were sacrificed under deep anesthesia with pentobarbital sodium (50 mg/kg body weight) and transcardiacly perfused with phosphate-buffered saline (PBS, pH 7.4). The brains were subsequently removed from the skull. The left hemisphere brain tissue was saved for western blotting and stored at -80 °C. The right hemisphere of the mouse brain tissue was fixed in 4% paraformaldehyde in 1 × PBS (pH 7.4) for 24 h and stored in 1 × PBS (pH 7.4) at 4 °C ^26^. For immunofluorescence staining, the fixed right brain hemisphere tissues were dehydrated via a vacuum infiltration processor (Leica ASP200S, Germany) and embedded in paraffin via an Arcadia H heated embedding workstation (Leica, Germany).

### 2.7 Protein extraction and western blotting

The cortices and hippocampi of the mice (n = 4 in each group) were lysed in RIPA buffer supplemented with a 0.1% protease inhibitor cocktail on ice. The protein concentration was measured with a Bradford kit (Bio-Rad). The proteins were analyzed by Western blotting as described previously ^27, 28^. The lysates were separated on a 7.5–15% SDS–polyacrylamide gel electrophoresis (PAGE) gel, and the proteins were transferred onto 0.22/0.45 μm polyvinylidene difluoride (PVDF) membranes. After the membranes were blocked with 5% milk, they were incubated with primary antibodies (**Supplementary Table 1**) at 4 °C overnight. The PVDF membranes were washed and then incubated with secondary peroxidase-coupled anti-mouse or anti-rabbit antibodies (1:5000) at room temperature for 1 h. The immunoreactive bands were visualized with Immobilon Western Horseradish peroxidase substrate luminol reagent (Millipore) via a ChemiDoc™ MP imaging system (Bio-Rad, USA).

### 2.8 Immunofluorescence staining and confocal imaging

Coronal sections of the mouse brains were cut at a thickness of 6 μm via a microtome (Leica RM2245, Germany). Dewaxed and rehydrated hippocampal sections were blocked in TBST buffer (tris-buffered saline (TBS) with Tween 20) with 5% bovine serum albumin for 1 h and then incubated with primary antibody 15A3 against 8-hydroxyguanosine (8-OHdG), 8-hydroxyguanosine (8-OHG) and the primary antibody NeuN at 4°C overnight ^29^. After being washed, the sections were incubated with AlexaFluor488-conjugated anti-mouse IgGs (1:200, Invitrogen, USA) for 1 h. After being washed with TBS, the sections and coverslips were mounted with vector anti-fading mounting medium (Vector Laboratories, Burlingame, CA, USA). The fluorescence intensity was imaged via a Leica SP8 confocal microscope at 100× magnification. For each cell, the total area corresponding to 15A3- and NeuN-related fluorescence was quantified via ImageJ 1.49 V software (NIH, US). The experiments were performed on 8 sections.

### 2.9 Statistical analysis

Statistical analysis was performed via SPSS software (version 23.0) or GraphPad Prism 10.0 software. The results are presented as the means ± SD. The Morris water maze behavior data were assessed via two-way repeated-measures ANOVA. The other parameters were analyzed via one-way ANOVA followed by Tukey’s *post hoc* test for multiple comparisons. Significance was set at p < 0.05.

## Results

### 3.1. Network pharmacology analysis and molecular docking for ISL

First, we obtained the intersection of drug-target genes and AD-related target genes for ISL. A set of 174 intersecting genes related to both AD and drugs was identified (S**Fig. 1**). These 174 intersecting target genes were subsequently imported into the STRING database (https://string-db.org/) for the prediction of protein_protein interactions. After topological analysis of the network, degree values were used to determine the size and color of the nodes; while combined score values were used to represent the thickness of the edges. The protein_protein interaction network (PPI network) for the core targets was constructed via Cytoscape 3.9.1 (**SFig. 1**). KEGG pathway analysis was performed via the DAVID database. A total of 126 pathways were screened, with 79 pathways related to ISL-associated diseases selected on the basis of p < 0.01. The top 30 KEGG metabolic pathways were visualized in a bubble chart based on P values (**SFig. 1**).

Molecular docking analysis was conducted for two selected targets mTOR and ERK2 with ISL (**Fig. 1**). 3D and 2D images of ISL binding to mTOR, with a binding energy of -7.1 kcal/mol. The amino acids of mTOR that form hydrogen bonds with ISL were Asp911, Asn1899, Thr1934, and Asn2147. The hydrogen bond lengths were 1.7 Å, 1.86 Å, 2.21 Å, and 2.02 Å, respectively (**Fig. 1A, B**). 3D and 2D images of ISL binding to ERK2, with a binding energy of -8.2 kcal/mol, indicating favorable binding interactions; the amino acids of ERK2 that form hydrogen bonds with ISL are Asp106, Met108, Ser153, and Asp167. The hydrogen bond lengths are 2.19 Å, 1.86 Å, 1.97 Å, and 1.72 Å, respectively (**Fig. 1C, D**). These residues may serve as key residues for the interaction between small molecules and proteins.

### 3.2 ISL rescued impaired learning and memory in STZ-induced model mice

Next, we assessed the effect of ISL on ICV-induced alterations in the behavior of the mice (**Fig. 1C-E**). ICV injection was confirmed by Trypan blue staining of coronal brain slices from additional mice that were injected in each group (**Fig. 2A**). The Morris water maze method was used to assess changes in learning and memory ability. After 4 days of learning, the last day was designated for testing the learning outcomes of the mice. On the fifth day, compared with the control group, the STZ group presented an increase in the time to first reach the platform quadrant (p=0.0122; **Fig. 2F**); a decrease in the number of platform crossings (p<0.0001; **Fig. 2G**), the time spent in the target quadrant (p=0.0022; **Fig. 2H**), and the distance traveled in the target quadrant (p=0.0004; **Fig. 2I**); compared with the STZ group, the STZ+ISL group presented a decrease in the time to first reach the platform quadrant (p=0.0305; **Fig. 2F**); and an increase in the number of platform crossings (p=0.0033; **Fig. 2G**), the time spent in the target quadrant (p=0.0368; **Fig. 2H**), and the distance traveled in the target quadrant (p=0.0043; **Fig. 2I**).

**Fig. 2.**
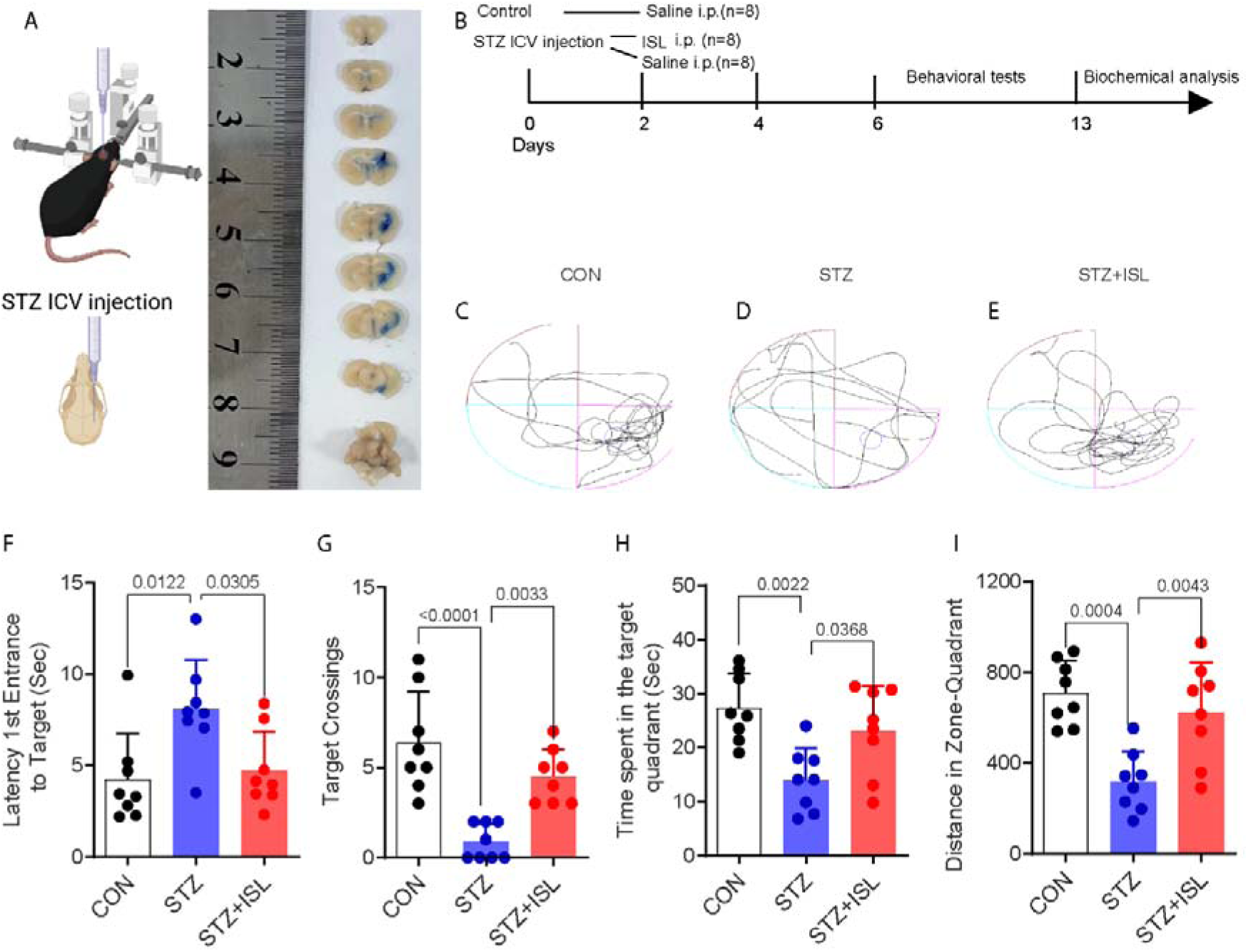
ISL treatment (i.p.) ameliorated STZ-induced cognitive impairment in mice. A) Scheme of the ICV injection of STZ (medial/lateral (ML): 2.5 mm, anterior/posterior (AP): -1.5 mm) in C57BL/6 mice and Trypan blue staining of the coronal brain slices. B) Experimental timeline for the ISL treatment and behavior test. C_E) Track of swimming; F) Time latency from the 1^st^ entrance to the target (second); G) Number of times the mice crossed the platform. H) Time spent by the mice in the target quadrant. I) Distance traveled by the mice in the target quadrant (n=8/per group).

### 3.3 ISL ameliorates tau hyperphosphorylation, the mTOR/ERK1/2 pathway and the upregulation of the GSK-3**β** pathway to prevent toxicity in STZ-induced mice

Next, we assessed the impact of ISL on STZ-induced tau phosphorylation in the cortex and hippocampus of mice injected with STZ (**Fig. 3**). We observed no changes in total tau protein (tau5) levels across all groups in the cortex or hippocampus of the mice in the control, STZ, and STZ+ISL groups (**Fig. 3C, G**). Compared with the control treatment, STZ treatment led to increased phosphorylation at Ser396 in the cortex and hippocampus of the mice (300% p<0.0001, 40% p<0.0001; **Fig. 3B, F**). Compared with the control treatment, STZ treatment led to increased p-tauSer396/tau5 in the cortex and hippocampus of the mice (%, 300% p<0.0001, 40% p<0.0001; **Fig. 3D, H**). Treatment with ISL partially reduced tauSer396 phosphorylation in the cortex (p=0.0029) but not in the hippocampus (p=0.0004) in the STZ-induced model mice (**Fig. 3B, F**). Compared with the control, ISL partially reduced the p-tauSer396/tau5 ratio in the cortex (%, p<0.0001) and completely reduced the p-tauSer396/tau5 ratio in the hippocampus of the STZ-induced model mice (p<0.0001, **Fig. 3D, H**).

**Fig. 3.**
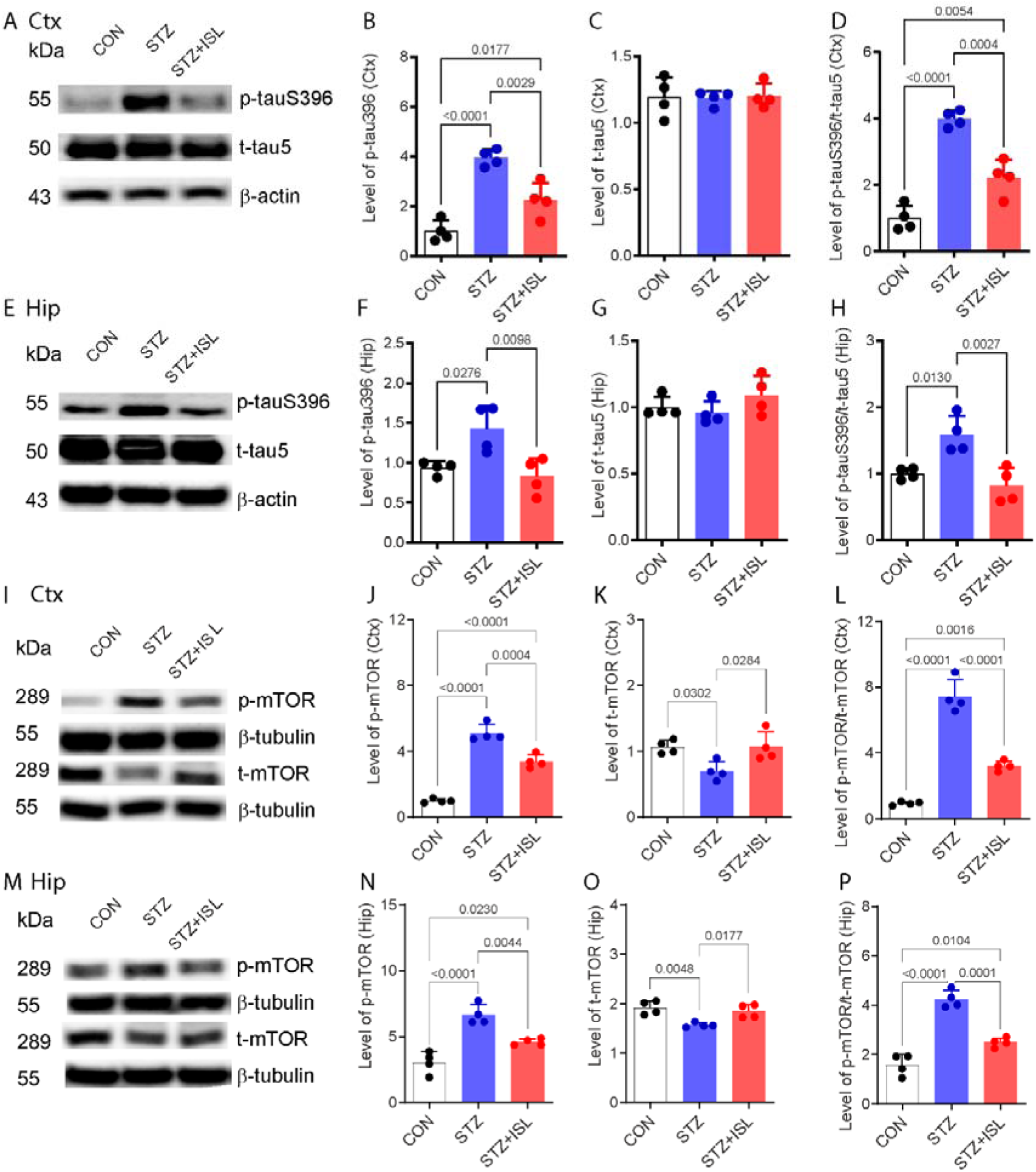
ISL treatment (i.p.) reversed the STZ-induced increase in TauS396 and mTOR phosphorylation in the mouse brain. A-H) Western blot analysis of tau proteins in the cortex and hippocampus. B, F) Levels of p-tauSer396. C, G) t-tau5 normalized to the control. D, H) p-TauSer396/t-tau5 ratio (n=4/per group). I_P) Western blot analysis of the protein levels of mTOR in the cortex and hippocampus. J, N) Levels of p-mTORS2448 and K, O) t-mTOR normalized to those of the control; L, P) p-mTORS2448/t-mTOR ratio (n=4/per group).

Next, we examined the effects on the expression and phosphorylation of mTOR, ERK1/2, and GSK-3β in each group (**Figs. 3, 4**). Compared with the control group, STZ treatment led to reduced t-mTOR in the cortex and hippocampus of the mice in the STZ group (40% p=0.0302, 20% p=0.0048). Treatment with ISL partially led to increased t-mTOR in the cortex and hippocampus of the STZ-induced mice (p=0.0284, p=0.0177). Compared with the control group, STZ treatment led to increased p-mTOR in the cortex and hippocampus (400% p<0.0001, 100% p<0.0001), as did the p-mTOR/t-mTOR ratio in the cortex and hippocampus of the mice in the STZ group (600% p<0.0001, 130% p<0.0001). Treatment with ISL partially reduced p-mTOR in the cortex and hippocampus (p=0.0004, p=0.0044). and p-mTOR/t-mTOR ratios in the cortex and hippocampus of the STZ-induced mice (p<0.0001, p<0.0001) (**Fig. 3J-L**).

**Fig. 4.**
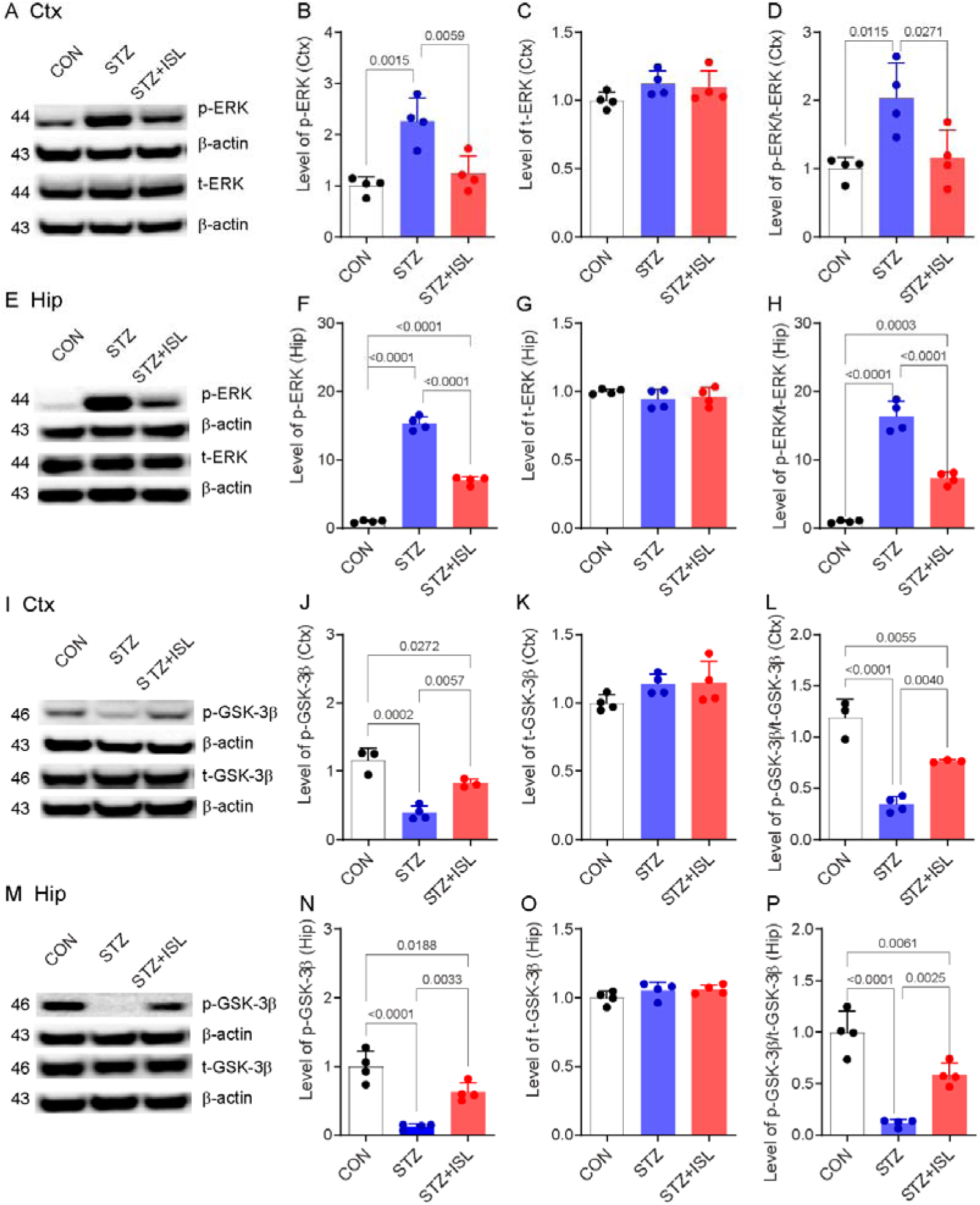
ISL treatment (i.p.) alleviated STZ-induced alterations in ERK and GSK-3β phosphorylation in the mouse brain. A-H) Western blot analysis of the protein levels of ERK1/2 in the cortex and hippocampus. B, F) Levels of p-ERK (T202/Y204) and C, G) t-ERK normalized to those of the control. D, H) p-ERK (T202/Y204)/t-ERK ratio (n=4/per group). I_P) Western blot analysis of the protein levels of GSK-3β in the cortex and hippocampus. J, N) Levels of p-GSK-3βSer9 and K, O) t-GSK-3β normalized to those of the control; L, P) p-GSK-3βSer9/t-GSK-3β ratio (n=4/per group).

We observed no changes in t-ERK1/2 protein levels in the cortex or in the hippocampus across all groups. Compared with the control group, STZ treatment led to increased p-ERK1/2 (T202/Y204) in the cortex and hippocampus (approximately 110% p=0.0015, 1400% p<0.0001) and increased p-ERK1/2 (T202/Y204)/t-ERK1/2 in the cortex and hippocampus of the mice in the STZ group (approximately 110% p<0.0001, 1400% p<0.0001). ISL treatment completely restored the p-ERK1/2 (T202/Y204) level in the cortex and hippocampus (p=0.0059, p<0.0001), as did the p-ERK1/2 (T202/Y204)/t-ERK1/2 ratio in the cortex and hippocampus of the STZ-induced model mice (p=0.0271, p<0.0001) (**Fig. 4A-H**).

In addition, we observed no changes in t-GSK-3β protein levels in the cortex or in the hippocampus across all groups. Compared with the control group, STZ treatment led to reduced p-GSK-3β (Ser9) in the cortex and hippocampus (approximately 60%, p=0.0002; 80%, p<0.0001) and p-GSK-3β (Ser9)/t-GSK-3β in the cortex and hippocampus of the mice in the STZ group (70%, p<0.0001; 85%, p<0.0001). ISL treatment partially (by half) decreased the p-GSK-3β (Ser9) level in the cortex and hippocampus (p=0.0057, p=0.0033) and the p-GSK-3β (Ser9)/t-GSK-3β level in the cortex and hippocampus of the STZ-induced model mice (p=0.0040, p=0.0025) (**Fig. 4I-P**). These findings suggest that the effects of ISL are associated with the deactivation of mTOR and ERK1/2 (T202/Y204) and the activation of GSK-3β, indicating a defensive response against oxidative stress

### 3.4 ISL ameliorated oxidative stress-induced mitochondrial impairment in STZ-induced model mice

To study oxidative stress, we used 15A3 antibodies targeting 8-OHdG and 8-OHG, which are indicators of DNA and RNA damage. The intensity of oxidative marker 15A3-positive staining was approximately 100% greater in the CA3 region of the mice in the STZ group than in those in the control group (p=0.0018). Treatment with ISL completely reversed the change in the intensity of the oxidative marker 15A3 in the CA3 region of the mice in the STZ+ISL group (p=0.0010, **Fig. 5A-B**).

**Fig. 5.**
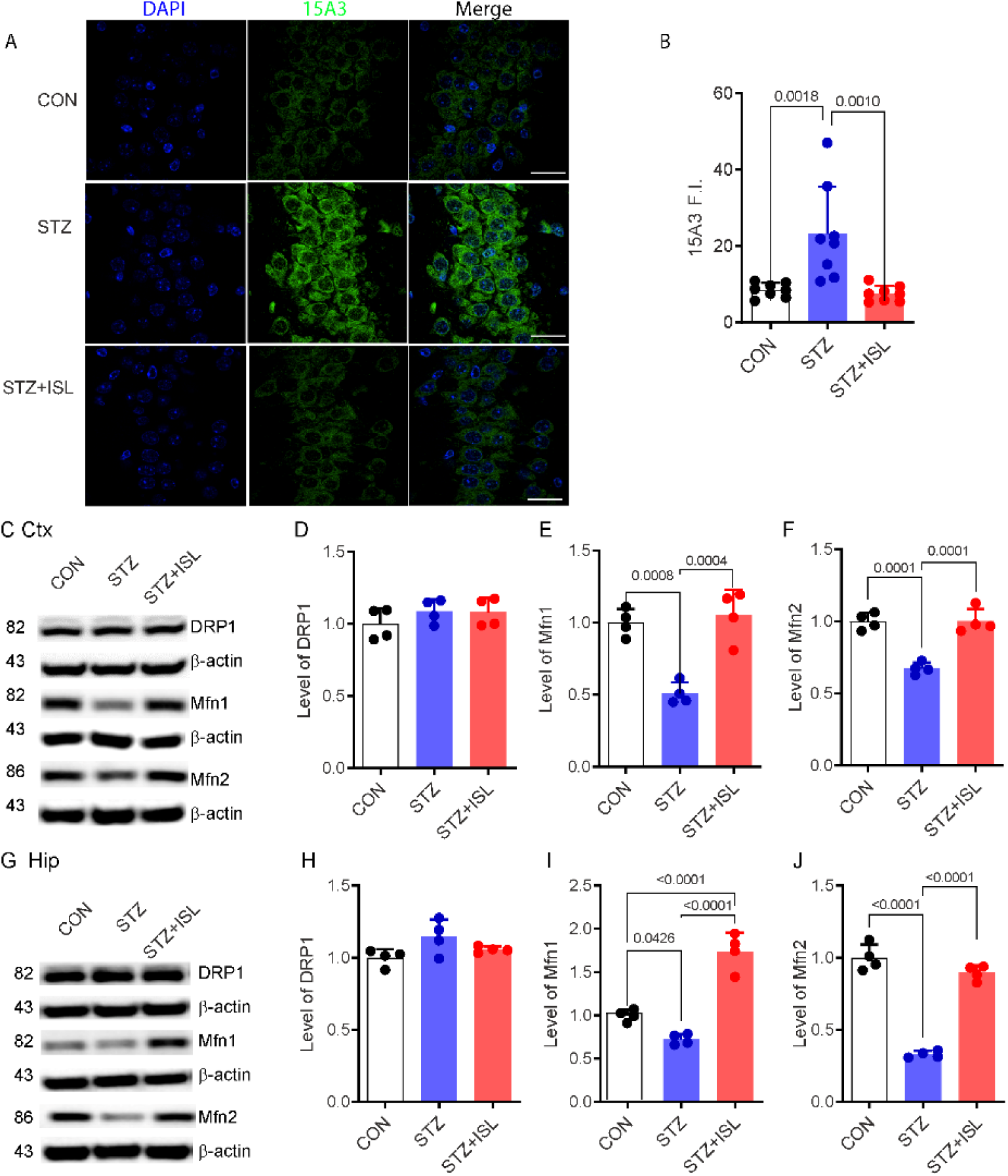
ISL treatment (i.p.) reversed the STZ-induced increase in oxidation indices and mitochondrial dysfunction in the mouse brain. A) 15A3 immunofluorescence staining of the CA3 area of the hippocampus. Nuclei were counterstained with DAPI (blue) and 15A3 (green); scale bar =10 μm; n=8 mice per group. B) Quantification of 15A3 fluorescence intensity. (C-H) Mitochondrial fission and fusion protein expression in the hippocampus and cortex. Western blot analysis of (D, I) Drp1, (F, J) Mfn1, and (G, K) Mfn2 expression. n=4 mice per group.

Next, we examined the effects of ISL treatment on the changes in mitochondrial function induced by STZ in mice. We observed that there were no changes in the protein levels of the mitochondrial fusion protein DRP1 in the cortex or in the hippocampus across all groups. Compared with those in the control group, the levels of the mitochondrial fission proteins Mfn1 and Mfn2 were lower in the cortex (50% p=0.0008, 40% p=0.0001) and hippocampus (30% p=0.0428, 70% p<0.0001) in the STZ group. ISL treatment completely reversed the changes in the levels of Mfn1 and Mfn2 in the cortex (p=0.0004, p=0.0001) and hippocampus (p<0.0001, p<0.0001) of the mice in the STZ group (**Fig. 5C-J**).

### 3.5 ISL rescued hippocampal neuronal loss and synaptic impairment in STZ-induced model mice

We subsequently assessed the impact of ISL treatment on STZ-induced synaptic and neuronal damage in mice. The intensity of neuronal marker NeuN-positive staining was approximately 20% lower in the CA3 region of the mice in the STZ group than in those in the control group (p<0.0001). Treatment with ISL mitigated the reduction in the intensity of NeuN in the CA3 region of the mice in the STZ+ISL group (p<0.0001) (**Fig. 6A-B**).

**Fig. 6.**
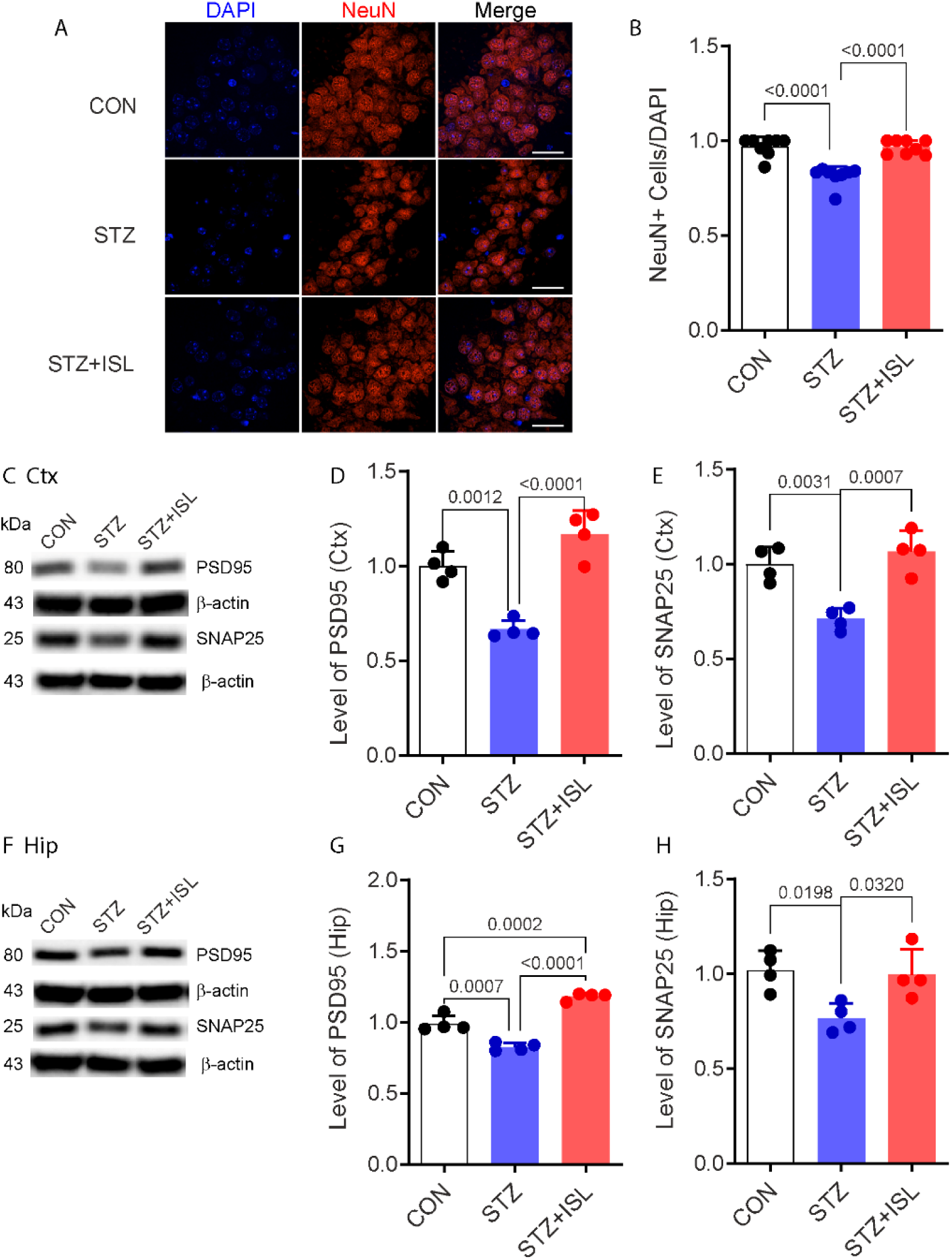
ISL treatment (i.p.) reversed STZ-induced synaptic protein and neuronal loss in the mouse brain. A) NeuN immunofluorescence staining in the CA3 area of the hippocampus. Nuclei were counterstained with DAPI (blue) and NeuN (red); scale bar =10 μm; n=8 mice per group. B) Quantification of NeuN-positive cells/cell nuclei. (C-H) Synaptic protein expression in the hippocampus and cortex. Western blot analysis of PSD95 (D, G) and SNAP25 (E, H) expression. n=4 mice per group.

Compared with those in the control group, the levels of the postsynaptic marker PSD95 and presynaptic marker SNAP25 were lower in the cortex (40% p=0.0012, 30% p=0.0031) and in the hippocampus (20% p=0.0007, 25% p=0.0198) in the STZ group. ISL treatment completely alleviated the STZ-induced reduction in PSD95 and SNAP25 levels in the cortex (p<0.0001, p=0.0007) and hippocampus (p<0.0001, p=0.0320) of the mice in the STZ group. (**Fig. 6C-H**).

## Discussion

Our data revealed that ISL treatment (i.p.) ameliorated tau hyperphosphorylation, oxidative stress, and mitochondrial and synaptic impairment in an STZ-induced mouse model via the mTOR, ERK1/2 and GSK-3β pathways. The interaction between ISL with mTOR, ERK was further supported by the molecular docking results. Moreover, ISL treatment reversed spatial learning deficits in the ICV-induced STZ-induced mouse model.

Here, we found that ISL reduced the increased levels of p-tauS396 in STZ-induced model mice. An earlier study showed that liquiritigenin had inhibitory effects on tau fibrillation and related neurotoxicity in a model of AD ^30^. This finding is in line with a previous study showing that phosphorylated tau was increased in both STZ-induced rat ^31^ and mouse models ^32^. The hyperphosphorylation of tau is a complex and multifaceted process that is central to the development of neurofibrillary tangles in the brains of individuals with AD. Phosphorylation at Ser396--404 is recognized as one of the early events in the progression of AD ^33^. The phosphorylation of tau at Ser396 plays a pivotal role in the hyperphosphorylation process and the subsequent formation of the tau tangle ^34^. Phosphorylation at Ser396 can decrease the ability of tau to regulate microtubule dynamic stability ^35^ and lead to a pathological tau conformation that is more prone to aggregation ^36^. Currently, two tau antibodies are in clinical trials: the C10.2 antibody specifically targets tau phosphorylated at Ser396 ^37^, and the liposome-based vaccine ACI-35 has a C-terminal tau fragment (393–408 residues).

Here, we found that ICV injection of STZ in the mouse brain induced the phosphorylation of mTOR (S2448) and ERK1/2 (T202/Y204) and that ISL ameliorated the phosphorylation of both mTOR (S2448) and ERK1/2 (T202/Y204) in STZ-injected mice. mTOR and ERK1/2 are among the most important serine/threonine kinases in eukaryotic cells, play prominent roles in tau hyperphosphorylation, and are therapeutic targets for AD ^38^. ERK1/2 signaling is involved in the modulation of synaptic plasticity and neuronal survival, which are compromised in AD ^39^. Abnormal activation of ERK1/2 has been observed in AD and is associated with increased tau phosphorylation and neuronal apoptosis. In line with our observations, increased Sirt2-induced tau phosphorylation through ERK activation has been reported in a STZ-induced AD-like mouse model ^32^. The mTOR signaling pathway is a key regulator of protein synthesis and cell growth and is an important target in AD. Dysregulation of mTOR signaling can lead to imbalanced tau phosphorylation through its effects on protein phosphatase 2 and GSK-3β activity ^40^. We previously reported that increased expression of p-mTOR (S2448) and p-ERK1/2 (T389) in postmortem AD brains was associated with the accumulation of hyperphosphorylated tau ^41^. An earlier study showed that mTOR inhibition of mTORC1 improved STZ-induced AD-like impairments in mice ^9^. In addition, the mTOR-mediated hyperphosphorylation of tau in the hippocampus is involved in cognitive deficits in STZ-induced diabetic ^42^ model mice ^43^ ISL has previously been shown to alleviate diabetic symptoms by activating AMPK and inhibiting mTORC1 signaling in diet-induced diabetic mice ^44^. Pathways such as the nuclear factor erythroid 2-related factor 2 (NRF2)-related pathway, Nrf2/NF-κB, the Nrf2/heme oxygenase-1/SLC7a11/glutathione peroxidase 4 axis and the Nrf2/antioxidant response element (ARE) signaling pathway have also been shown to be involved in the effect of ISL ^17, 21, 45^.

Our data revealed that t-GSK-3β levels were similar between ISL-treated and untreated STZ-induced mice and that the levels of GSK-3β phosphorylated at Ser9 were significantly greater in the STZ-induced mice treated with ISL than in the STZ-induced mice. GSK-3β is a proline-directed serine/threonine kinase that plays a vital role in the regulation of glycogen metabolism. GSK-3β plays a critical role in the AD brain and is involved in tau hyperphosphorylation. Moreover, GSK-3β is involved in neuroinflammation and the maintenance of neuronal plasticity ^46^. The downregulation of GSK-3 activity may involve phosphorylation at the S21 site for GSK-3α and at the S9 site for GSK-3β ^46^. Given that Ser9 is an inhibitory phospho-site, these results indicate that the effect of ISL involves downregulating the activity of GSK-3β, thereby increasing GSK-3 activity. Elevated levels of the active form of GSK-3β have been detected in the tangle-bearing neurons of AD patients ^47^. Both in vitro and in vivo studies have demonstrated that the upregulation of GSK-3β induces tau hyperphosphorylation and memory deficits, whereas the inhibition of GSK-3 attenuates tau hyperphosphorylation and improves synaptic function. Additionally, GSK-3β signals through mTORC1 to regulate nutrient-induced mitochondrial activity independently of mitochondrial biogenesis ^48^. The inhibition of GSK-3β has been shown to stimulate nutrient-induced mitochondrial activity in cultured human neurons, mitochondrial activity and oxygen consumption ^49^. In line with our findings, the protective effects of ISL have also been shown to be exerted through inhibiting GSK-3β activity through increasing the expression levels of p-GSK-3β, thereby disinhibiting Nrf2 and upregulating the gene expression of Nrf2-controlled antioxidant genes and the ERK/mitochondrion pathway ^50^, preventing mitochondrial fission ^15, 51, 52^. Thus, the inhibition of GSK-3β is considered a potential therapeutic target for neurodegenerative diseases.

Studies have shown that the brains of AD patients and experimental animal models are exposed to oxidative stress through high levels of oxidized proteins, lipid peroxidation and oxidative modifications in nuclear and mitochondrial DNA. The oxidized nucleosides 8-OHdG and 8-OHG are markers of oxidative damage to DNA and RNA, respectively. An earlier study revealed that the immunoreactivity of 15A3 (for both 8OHdG and 8OHG) was more prominent in the cytoplasm and less prominent in the nucleus in neurons ^53^. Our results revealed that the ICV injection of STZ significantly increased the expression of the nucleic peroxidation product 15A3, indicating that oxidative stress leads to the degeneration of neurons. Treatment with ISL reversed the STZ-induced increase in 15A3 (8-OHdG and 8-OHG) in the hippocampus of the STZ-induced mice. Previous in vitro studies have shown that ISL protects neuronal cells from 6-hydroxydopamine (6-OHDA)- and glutamate-induced neurotoxicity by reducing the production of ROS ^54^.

Mitochondria play key roles in connection with cell signaling and oxidative stress, are associated with mitochondrial dynamics and are impaired in models of AD ^27, 55^. Mitochondrial dysfunction is also detected by in vivo imaging in the parahippocampus in patients with early-stage AD ^56^. A disrupted mitochondrial response to nutrients has been shown to be a presymptomatic event in the brain of the APP^SAA^ knock-in mouse model of AD ^49^. Insulin deficiency and intranasal insulin alter brain mitochondrial function and are considered potential factors for dementia in individuals with diabetes ^57^. The balance of mitochondrial dynamics between fusion and fission influences the morphology and function of mitochondria. We observed elevated levels of the fusion proteins Mfn1 and Mfn2 in cortical and hippocampal neurons in the brains of STZ-induced model mice, with no changes in the levels of the mitochondrial fission protein Drp1. Treatment with ISL alleviated this reduction in Mfn1 and Mfn2 expression in the STZ-induced model mice. These findings suggest that STZ induced an imbalance in mitochondrial fission/fusion. Once the balance of fission/fusion is perturbed, the mitochondria undergo abnormal morphological and functional changes. A previous proteomics study demonstrated alterations in mitochondria, such as atp5f1d, atp5f1a, and uqcrc1, in an ICV-STZ-induced rat model ^58^. In line with our findings, ISL has been shown to reduce lipopolysaccharide-induced inflammation by preventing mitochondrial fission ^51^ and improving mitochondrial function in cellular models of oxidative and nitrosative stress ^59^ and neurodegenerative diseases ^11^. Additionally, reports indicate that ISL can improve mitochondrial function via the peroxisome proliferator-activated receptor gamma-dependent pathway and the Nrf2-mediated pathway ^11, 14, 60^.

An earlier study revealed that ISL protects against cognitive impairment and neuronal injury induced by the injection of lipopolysaccharides ^15^. Our results suggested that ISL treatment reversed spatial learning and memory impairments in the STZ-induced model mice. The spatial memory impairment we observed in the STZ-induced mice is in line with the previously reported deterioration of learning and memory in this model ^8^. Memory performance is associated with hippocampal morphology and function, and STZ reportedly causes the loss of pyramidal neurons in the CA1 region ^61^. Moreover, we showed that ISL treatment reversed the decreases in the expression levels of the presynaptic protein SNAP 25 and the postsynaptic protein PSD95 in both the cortex and hippocampus of the mice.

There are several limitations in this study. First, only male mice were included in the experimental design. Second, only the Morris water maze test was used to assess the spatial learning ability of the mice. The use of a panel of behavior tests will provide comprehensive insights into the effect of ISL on STZ-induced cognitive impairment. Third, we focused on alterations in the tau396, mTOR, ERK, and GSK-3β pathways in the present study, whereas many other pathways and pathophysiological changes, such as amyloid-beta and neuroinflammation, might be involved in the effect of ISL on STZ-induced nephropathy in mice. Moreover, treatment with ISL at different doses and via other routes of administration, such as oral intake, remains to be investigated, as the efficacy and specific outcomes can vary on the basis of factors such as animal strain, age, and dosage ^62^. A longitudinal study using in vivo imaging ^63^ of the treatment effect of ISL on tau, neuroinflammation, and mitochondrial dysfunction in an animal model will provide further systematic insights.

## Conclusions

In conclusion, treatment with ISL (i.p.) attenuated p-tauS396 levels and mTOR and ERK activity; increased GSK-3β activity; and alleviated synaptic impairment, oxidative stress and cognitive deficits in an ICV-STZ-injected mouse model.

## Ethics approval and consent to participate

All experimental protocols were approved by the Guiyang Regional Animal Care Center and Ethics Committee (No. 2304628). All the experiments followed ARRIVE guidelines 2.0.

Our study did not require consent to participate, as there was no sample or data from humans.

## Consent for publication

Not relevant

## Author Contributions

ZT contributed to the study conception and design. ST, and YTD performed the experiments. YX and YSW contributed to the data collection and data analysis. RN and ZT interpreted the data. RN, ST and ZT wrote the manuscript. ZT and XLQ acquired funding. All authors approved the manuscript before submission.

## Availability of data and material

The data are available upon reasonable request.

## References

(1) Scheltens, P.; De Strooper, B.; Kivipelto, M.; Holstege, H.; Chételat, G.; Teunissen, C. E.; Cummings, J.; van der Flier, W. M. Alzheimer’s disease. The Lancet 2021, 397 (10284), 1577–1590. DOI: 10.1016/S0140-6736(20)32205-4 (acccessed 2021/09/04).

(2) Islam, M. T. Oxidative stress and mitochondrial dysfunction-linked neurodegenerative disorders. Neurological research 2017, 39 (1), 73–82. DOI: 10.1080/01616412.2016.1251711 From NLM. Niedzielska, E.; Smaga, I.; Gawlik, M.; Moniczewski, A.; Stankowicz, P.; Pera, J.; Filip, M. Oxidative Stress in Neurodegenerative Diseases. Molecular neurobiology 2016, 53 (6), 4094–4125. DOI: 10.1007/s12035-015-9337-5 From NLM. Tönnies, E.; Trushina, E. Oxidative Stress, Synaptic Dysfunction, and Alzheimer’s Disease. Journal of Alzheimer’s disease : JAD 2017, 57 (4), 1105–1121. DOI: 10.3233/jad-161088 From NLM.

(3) Wang, J.; Huang, Q.; He, K.; Li, J.; Guo, T.; Yang, Y.; Lin, Z.; Li, S.; Vanderlinden, G.; Huang, Y.;, et al. Presynaptic density determined by SV2A PET is closely associated with postsynaptic metabotropic glutamate receptor 5 availability and independent of amyloid pathology in early cognitive impairment. Alzheimers Dement 2024. DOI: 10.1002/alz.13817 From NLM.

(4) Kong, Y.; Cao, L.; Wang, J.; Zhuang, J.; Xie, F.; Zuo, C.; Huang, Q.; Shi, K.; Rominger, A.; Li, M.;, et al. In vivo reactive astrocyte imaging using [18F]SMBT-1 in tauopathy and familial Alzheimer’s disease mouse models: A multi-tracer study. Journal of the Neurological Sciences 2024, 123079. DOI: 10.1016/j.jns.2024.123079. Kong, Y.; Maschio, C. A.; Shi, X.; Xie, F.; Zuo, C.; Konietzko, U.; Shi, K.; Rominger, A.; Xiao, J.; Huang, Q.; et al. Relationship Between Reactive Astrocytes, by [(18)F]SMBT-1 Imaging, with Amyloid-Beta, Tau, Glucose Metabolism, and TSPO in Mouse Models of Alzheimer’s Disease. Mol Neurobiol 2024. DOI: 10.1007/s12035-024-04106-7 From NLM. Park, L.; Hochrainer, K.; Hattori, Y.; Ahn, S. J.; Anfray, A.; Wang, G.; Uekawa, K.; Seo, J.; Palfini, V.; Blanco, I.; et al. Tau induces PSD95-neuronal NOS uncoupling and neurovascular dysfunction independent of neurodegeneration. Nat Neurosci 2020, 23 (9), 1079–1089. DOI: 10.1038/s41593-020-0686-7 From NLM.

(5) Minhas, P. S.; Jones, J. R.; Latif-Hernandez, A.; Sugiura, Y.; Durairaj, A. S.; Wang, Q.; Mhatre, S. D.; Uenaka, T.; Crapser, J.; Conley, T.;, et al. Restoring hippocampal glucose metabolism rescues cognition across Alzheimer’s disease pathologies. Science 2024, 385 (6711), eabm6131. DOI: 10.1126/science.abm6131 From NLM.

(6) Grieb, P. Intracerebroventricular Streptozotocin Injections as a Model of Alzheimer’s Disease: in Search of a Relevant Mechanism. Molecular Neurobiology 2016, 53 (3), 1741–1752. DOI: 10.1007/s12035-015-9132-3. Salkovic-Petrisic, M.; Knezovic, A.; Hoyer, S.; Riederer, P. What have we learned from the streptozotocin-induced animal model of sporadic Alzheimer’s disease, about the therapeutic strategies in Alzheimer’s research. J Neural Transm (Vienna) 2013, 120 (1), 233–252. DOI: 10.1007/s00702-012-0877-9 From NLM. Zhang, J.; Liu, L.; Zhang, Y.; Yuan, Y.; Miao, Z.; Lu, K.; Zhang, X.; Ni, R.; Zhang, H.; Zhao, Y.; et al. ChemR23 signaling ameliorates cognitive impairments in diabetic mice via dampening oxidative stress and NLRP3 inflammasome activation. Redox Biology 2022, 58, 102554. DOI: https://doi.org/10.1016/j.redox.2022.102554.

(7) Rostami, F.; Javan, M.; Moghimi, A.; Haddad-Mashadrizeh, A.; Fereidoni, M. Streptozotocin-induced hippocampal astrogliosis and insulin signaling malfunction as experimental scales for subclinical sporadic Alzheimer model. Life Sciences 2017, 188, 172–185. DOI: 10.1016/j.lfs.2017.08.025. Silva, S. S. L.; Tureck, L. V.; Souza, L. C.; Mello-Hortega, J. V.; Piumbini, A. L.; Teixeira, M. D.; Furtado-Alle, L.; Vital, M. A. B. F.; Souza, R. L. R. Animal model of Alzheimer’s disease induced by streptozotocin: New insights about cholinergic pathway. Brain Research 2023, 1799, 148175. DOI: https://doi.org/10.1016/j.brainres.2022.148175. Kamat, P. K. Streptozotocin induced Alzheimer’s disease like changes and the underlying neural degeneration and regeneration mechanism. Neural Regen Res 2015, 10 (7), 1050–1052. DOI: 10.4103/1673-5374.160076 From NLM. Gáspár, A.; Hutka, B.; Ernyey, A. J.; Tajti, B. T.; Varga, B. T.; Zádori, Z. S.; Gyertyán, I. Performance of the intracerebroventricularly injected streptozotocin Alzheimer’s disease model in a translationally relevant, aged and experienced rat population. Scientific Reports 2022, 12 (1), 20247. DOI: 10.1038/s41598-022-24292-5. Dos Santos, J. P. A.; Vizuete, A.; Hansen, F.; Biasibetti, R.; Gonçalves, C. A. Early and Persistent O-GlcNAc Protein Modification in the Streptozotocin Model of Alzheimer’s Disease. J Alzheimers Dis 2018, 61 (1), 237–249. DOI: 10.3233/jad-170211 From NLM. Kelliny, S.; Lin, L.; Deng, I.; Xiong, J.; Zhou, F.; Al-Hawwas, M.; Bobrovskaya, L.; Zhou, X. F. A New Approach to Model Sporadic Alzheimer’s Disease by Intracerebroventricular Streptozotocin Injection in APP/PS1 Mice. Mol Neurobiol 2021, 58 (8), 3692–3711. DOI: 10.1007/s12035-021-02338-5 From NLM.

(8) Fan, M.; Liu, S.; Sun, H. M.; Ma, M. D.; Gao, Y. J.; Qi, C. C.; Xia, Q. R.; Ge, J. F. Bilateral intracerebroventricular injection of streptozotocin induces AD-like behavioral impairments and neuropathological features in mice: Involved with the fundamental role of neuroinflammation. Biomed Pharmacother 2022, 153, 113375. DOI: 10.1016/j.biopha.2022.113375 From NLM.

(9) Cao, Y.; Liu, B.; Xu, W.; Wang, L.; Shi, F.; Li, N.; Lei, Y.; Wang, J.; Tian, Q.; Zhou, X. Inhibition of mTORC1 improves STZ-induced AD-like impairments in mice. Brain Res Bull 2020, 162, 166–179. DOI: 10.1016/j.brainresbull.2020.06.002 From NLM.

(10) Honda, H.; Nagai, Y.; Matsunaga, T.; Okamoto, N.; Watanabe, Y.; Tsuneyama, K.; Hayashi, H.; Fujii, I.; Ikutani, M.; Hirai, Y.;, et al. Isoliquiritigenin is a potent inhibitor of NLRP3 inflammasome activation and diet-induced adipose tissue inflammation. J Leukoc Biol 2014, 96 (6), 1087–1100. DOI: 10.1189/jlb.3A0114-005RR From NLM.

(11) Denzer, I.; Münch, G.; Friedland, K. Modulation of mitochondrial dysfunction in neurodegenerative diseases via activation of nuclear factor erythroid-2-related factor 2 by food-derived compounds. Pharmacol Res 2016, 103, 80–94. DOI: 10.1016/j.phrs.2015.11.019 From NLM.

(12) Sharma, R.; Singla, R. K.; Banerjee, S. Revisiting Licorice as a functional food in the management of neurological disorders: Bench to trend. Neurosci Biobehav Rev 2023, 155, 105452. DOI: 10.1016/j.neubiorev.2023.105452 From NLM.

(13) Yokoyama, T.; Hisatomi, K.; Oshima, S.; Tanaka, I.; Okada, T.; Toyooka, N. Discovery and optimization of isoliquiritigenin as a death-associated protein kinase 1 inhibitor. Eur J Med Chem 2024, 279, 116836. DOI: 10.1016/j.ejmech.2024.116836 From NLM. Gay, N. H.; Suwanjang, W.; Ruankham, W.; Songtawee, N.; Wongchitrat, P.; Prachayasittikul, V.; Prachayasittikul, S.; Phopin, K. Butein, isoliquiritigenin, and scopoletin attenuate neurodegeneration via antioxidant enzymes and SIRT1/ADAM10 signaling pathway. RSC Adv 2020, 10 (28), 16593–16606. DOI: 10.1039/c9ra06056a From NLM. Yang, E.-J.; Min, J. S.; Ku, H.-Y.; Choi, H.-S.; Park, M.-k.; Kim, M. K.; Song, K.-S.; Lee, D.-S. Isoliquiritigenin isolated from Glycyrrhiza uralensis protects neuronal cells against glutamate-induced mitochondrial dysfunction. Biochemical and Biophysical Research Communications 2012, 421 (4), 658–664. DOI: https://doi.org/10.1016/j.bbrc.2012.04.053.

(14) Shi, D.; Yang, J.; Jiang, Y.; Wen, L.; Wang, Z.; Yang, B. The antioxidant activity and neuroprotective mechanism of isoliquiritigenin. Free Radic Biol Med 2020, 152, 207–215. DOI: 10.1016/j.freeradbiomed.2020.03.016 From NLM.

(15) Zhu, X.; Liu, J.; Chen, S.; Xue, J.; Huang, S.; Wang, Y.; Chen, O. Isoliquiritigenin attenuates lipopolysaccharide-induced cognitive impairment through antioxidant and anti-inflammatory activity. BMC Neurosci 2019, 20 (1), 41. DOI: 10.1186/s12868-019-0520-x From NLM.

(16) Long, Y.; Yang, Q.; Xiang, Y.; Zhang, Y.; Wan, J.; Liu, S.; Li, N.; Peng, W. Nose to brain drug delivery - A promising strategy for active components from herbal medicine for treating cerebral ischemia reperfusion. Pharmacol Res 2020, 159, 104795. DOI: 10.1016/j.phrs.2020.104795 From NLM.

(17) Lan, X.; Wang, Q.; Liu, Y.; You, Q.; Wei, W.; Zhu, C.; Hai, D.; Cai, Z.; Yu, J.; Zhang, J.;, et al. Isoliquiritigenin alleviates cerebral ischemia-reperfusion injury by reducing oxidative stress and ameliorating mitochondrial dysfunction via activating the Nrf2 pathway. Redox Biol 2024, 77, 103406. DOI: 10.1016/j.redox.2024.103406 From NLM.

(18) Zhu, X.; Liu, J.; Huang, S.; Zhu, W.; Wang, Y.; Chen, O.; Xue, J. Neuroprotective effects of isoliquiritigenin against cognitive impairment via suppression of synaptic dysfunction, neuronal injury, and neuroinflammation in rats with kainic acid-induced seizures. International Immunopharmacology 2019, 72, 358–366. DOI: 10.1016/j.intimp.2019.04.028.

(19) Liu, J. Q.; Zhao, X. T.; Qin, F. Y.; Zhou, J. W.; Ding, F.; Zhou, G.; Zhang, X. S.; Zhang, Z. H.; Li, Z. B. Isoliquiritigenin mitigates oxidative damage after subarachnoid hemorrhage in vivo and in vitro by regulating Nrf2-dependent Signaling Pathway via Targeting of SIRT1. Phytomedicine 2022, 105, 154262. DOI: 10.1016/j.phymed.2022.154262 From NLM.

(20) Huang, L.; Han, Y.; Zhou, Q.; Sun, Z.; Yan, J. Isoliquiritigenin attenuates neuroinflammation in mice model of Parkinson’s disease by promoting Nrf2/NQO-1 pathway. Transl Neurosci 2022, 13 (1), 301–308. DOI: 10.1515/tnsci-2022-0239 From NLM.

(21) Fu, Y.; Jia, J. Isoliquiritigenin Confers Neuroprotection and Alleviates Amyloid-β42-Induced Neuroinflammation in Microglia by Regulating the Nrf2/NF-κB Signaling. Front Neurosci 2021, 15, 638772. DOI: 10.3389/fnins.2021.638772 From NLM.

(22) Chen, Y. P.; Zhang, Z. Y.; Li, Y. P.; Li, D.; Huang, S. L.; Gu, L. Q.; Xu, J.; Huang, Z. S. Syntheses and evaluation of novel isoliquiritigenin derivatives as potential dual inhibitors for amyloid-beta aggregation and 5-lipoxygenase. Eur J Med Chem 2013, 66, 22–31. DOI: 10.1016/j.ejmech.2013.05.015 From NLM. Lee, H. K.; Yang, E. J.; Kim, J. Y.; Song, K. S.; Seong, Y. H. Inhibitory effects of Glycyrrhizae radix and its active component, isoliquiritigenin, on Aβ(25-35)-induced neurotoxicity in cultured rat cortical neurons. Arch Pharm Res 2012, 35 (5), 897–904. DOI: 10.1007/s12272-012-0515-y From NLM. Nair, A. C.; Benny, S.; Tp, A.; Sudheesh, M. S.; Lakshmi, P. K. Comprehensive profiling of traditional herbomineral formulation Manasamitra vatakam in rat brain following oral administration and in-silico screening of the identified compound for anti-Alzheimer’s activity. J Ethnopharmacol 2024, 119024. DOI: 10.1016/j.jep.2024.119024 From NLM.

(23) Wang, L.; Tang, Z.; Li, B.; Peng, Y.; Yang, X.; Xiao, Y.; Ni, R.; Qi, X. L. Myricetin ameliorates cognitive impairment in 3×Tg Alzheimer’s disease mice by regulating oxidative stress and tau hyperphosphorylation. Biomed Pharmacother 2024, 177, 116963. DOI: 10.1016/j.biopha.2024.116963 From NLM.

(24) Lai, C.; Chen, Z.; Ding, Y.; Chen, Q.; Su, S.; Liu, H.; Ni, R.; Tang, Z. Rapamycin Attenuated Zinc-Induced Tau Phosphorylation and Oxidative Stress in Rats: Involvement of Dual mTOR/p70S6K and Nrf2/HO-1 Pathways. Front Immunol 2022, 13, 782434. DOI: 10.3389/fimmu.2022.782434 From NLM.

(25) Vorhees, C. V.; Williams, M. T. Morris water maze: procedures for assessing spatial and related forms of learning and memory. Nature protocols 2006, 1 (2), 848–858. DOI: 10.1038/nprot.2006.116 PubMed.

(26) Ding, Y.; Liu, H.; Cen, M.; Tao, Y.; Lai, C.; Tang, Z. Rapamycin Ameliorates Cognitive Impairments and Alzheimer’s Disease-Like Pathology with Restoring Mitochondrial Abnormality in the Hippocampus of Streptozotocin-Induced Diabetic Mice. Neurochemical research 2021, 46 (2), 265–275. DOI: 10.1007/s11064-020-03160-6.

(27) Tang, Z.; Chen, Z.; Guo, M.; Peng, Y.; Xiao, Y.; Guan, Z.; Ni, R.; Qi, X. NRF2 Deficiency Promotes Ferroptosis of Astrocytes Mediated by Oxidative Stress in Alzheimer’s Disease. Molecular Neurobiology 2024. DOI: 10.1007/s12035-024-04023-9. Tang, Z.; Peng, Y.; Jiang, Y.; Wang, L.; Guo, M.; Chen, Z.; Luo, C.; Zhang, T.; Xiao, Y.; Ni, R.; et al. Gastrodin ameliorates synaptic impairment, mitochondrial dysfunction and oxidative stress in N2a/APP cells. Biochem Biophys Res Commun 2024, 719, 150127. DOI: 10.1016/j.bbrc.2024.150127 From NLM.

(28) Chen, Q.; Lai, C.; Chen, F.; Ding, Y.; Zhou, Y.; Su, S.; Ni, R.; Tang, Z. Emodin Protects SH-SY5Y Cells Against Zinc-Induced Synaptic Impairment and Oxidative Stress Through the ERK1/2 Pathway. Front Pharmacol 2022, 13, 821521. DOI: 10.3389/fphar.2022.821521 From NLM.

(29) Kecheliev, V.; Boss, L.; Maheshwari, U.; Konietzko, U.; Keller, A.; Razansky, D.; Nitsch, R. M.; Klohs, J.; Ni, R. Aquaporin 4 is differentially increased and dislocated in association with tau and amyloid-beta. Life Sciences 2023, 121593. DOI: 10.1016/j.lfs.2023.121593.

(30) Yuan, X.; Wang, Z.; Zhang, L.; Sui, R.; Khan, S. Exploring the inhibitory effects of liquiritigenin against tau fibrillation and related neurotoxicity as a model of preventive care in Alzheimer’s disease. Int J Biol Macromol 2021, 183, 1184–1190. DOI: 10.1016/j.ijbiomac.2021.05.041 From NLM.

(31) Du, L. L.; Chai, D. M.; Zhao, L. N.; Li, X. H.; Zhang, F. C.; Zhang, H. B.; Liu, L. B.; Wu, K.; Liu, R.; Wang, J. Z.;, et al. AMPK activation ameliorates Alzheimer’s disease-like pathology and spatial memory impairment in a streptozotocin-induced Alzheimer’s disease model in rats. J Alzheimers Dis 2015, 43 (3), 775–784. DOI: 10.3233/jad-140564 From NLM.

(32) Zhou, C.; Jung, C. G.; Kim, M. J.; Watanabe, A.; Abdelhamid, M.; Taslima, F.; Michikawa, M. Insulin Deficiency Increases Sirt2 Level in Streptozotocin-Treated Alzheimer’s Disease-Like Mouse Model: Increased Sirt2 Induces Tau Phosphorylation Through ERK Activation. Mol Neurobiol 2022, 59 (9), 5408–5425. DOI: 10.1007/s12035-022-02918-z From NLM.

(33) Xia, Y.; Prokop, S.; Giasson, B. I. "Don’t Phos Over Tau": recent developments in clinical biomarkers and therapies targeting tau phosphorylation in Alzheimer’s disease and other tauopathies. Mol Neurodegener 2021, 16 (1), 37. DOI: 10.1186/s13024-021-00460-5 From NLM. Neddens, J.; Temmel, M.; Flunkert, S.; Kerschbaumer, B.; Hoeller, C.; Loeffler, T.; Niederkofler, V.; Daum, G.; Attems, J.; Hutter-Paier, B. Phosphorylation of different tau sites during progression of Alzheimer’s disease. Acta Neuropathol Commun 2018, 6 (1), 52. DOI: 10.1186/s40478-018-0557-6 From NLM. Mondragón-Rodríguez, S.; Perry, G.; Luna-Muñoz, J.; Acevedo-Aquino, M. C.; Williams, S. Phosphorylation of tau protein at sites Ser(396-404) is one of the earliest events in Alzheimer’s disease and Down syndrome. Neuropathol Appl Neurobiol 2014, 40 (2), 121–135. DOI: 10.1111/nan.12084 From NLM.

(34) Wegmann, S.; Biernat, J.; Mandelkow, E. A current view on Tau protein phosphorylation in Alzheimer’s disease. Curr Opin Neurobiol 2021, 69, 131–138. DOI: 10.1016/j.conb.2021.03.003 From NLM.

(35) Kiris, E.; Ventimiglia, D.; Sargin, M. E.; Gaylord, M. R.; Altinok, A.; Rose, K.; Manjunath, B. S.; Jordan, M. A.; Wilson, L.; Feinstein, S. C. Combinatorial Tau pseudophosphorylation: markedly different regulatory effects on microtubule assembly and dynamic instability than the sum of the individual parts. J Biol Chem 2011, 286 (16), 14257–14270. DOI: 10.1074/jbc.M111.219311 From NLM.

(36) Jeganathan, S.; Hascher, A.; Chinnathambi, S.; Biernat, J.; Mandelkow, E. M.; Mandelkow, E. Proline-directed pseudo-phosphorylation at AT8 and PHF1 epitopes induces a compaction of the paperclip folding of Tau and generates a pathological (MC-1) conformation. J Biol Chem 2008, 283 (46), 32066–32076. DOI: 10.1074/jbc.M805300200 From NLM.

(37) Rosenqvist, N.; Asuni, A. A.; Andersson, C. R.; Christensen, S.; Daechsel, J. A.; Egebjerg, J.; Falsig, J.; Helboe, L.; Jul, P.; Kartberg, F.;, et al. Highly specific and selective anti-pS396-tau antibody C10.2 targets seeding-competent tau. Alzheimers Dement (N Y) 2018, 4, 521–534. DOI: 10.1016/j.trci.2018.09.005 From NLM.

(38) Griffin, R. J.; Moloney, A.; Kelliher, M.; Johnston, J. A.; Ravid, R.; Dockery, P.; O’Connor, R.; O’Neill, C. Activation of Akt/PKB, increased phosphorylation of Akt substrates and loss and altered distribution of Akt and PTEN are features of Alzheimer’s disease pathology. Journal of neurochemistry 2005, 93 (1), 105–117. DOI: 10.1111/j.1471-4159.2004.02949.x. Pei, J. J.; Hugon, J. mTOR-dependent signalling in Alzheimer’s disease. Journal of cellular and molecular medicine 2008, 12 (6B), 2525–2532. DOI: 10.1111/j.1582-4934.2008.00509.x. Liu, G. Y.; Sabatini, D. M. mTOR at the nexus of nutrition, growth, ageing and disease. Nature reviews. Molecular cell biology 2020, 21 (4), 183–203. DOI: 10.1038/s41580-019-0199-y From NLM. Yin, S.; Liu, L.; Gan, W. The Roles of Post-Translational Modifications on mTOR Signaling. International journal of molecular sciences 2021, 22 (4). DOI: 10.3390/ijms22041784 From NLM. Bockaert, J.; Marin, P. mTOR in Brain Physiology and Pathologies. Physiological reviews 2015, 95 (4), 1157–1187. DOI: https://doi.org/10.1152/physrev.00038.2014 From NLM.

(39) Davoody, S.; Asgari Taei, A.; Khodabakhsh, P.; Dargahi, L. mTOR signaling and Alzheimer’s disease: What we know and where we are? CNS Neurosci Ther 2024, 30 (4), e14463. DOI: 10.1111/cns.14463 From NLM. Chiang, G. G.; Abraham, R. T. Phosphorylation of Mammalian Target of Rapamycin (mTOR) at Ser-2448 IsMediated by p70S6 Kinase*. Journal of Biological Chemistry 2005, 280 (27), 25485–25490. DOI: https://doi.org/10.1074/jbc.M501707200.

(40) Caccamo, A.; Magrì, A.; Medina, D. X.; Wisely, E. V.; López-Aranda, M. F.; Silva, A. J.; Oddo, S. mTOR regulates tau phosphorylation and degradation: implications for Alzheimer’s disease and other tauopathies. Aging Cell 2013, 12 (3), 370–380. DOI: 10.1111/acel.12057 From NLM.

(41) An, W. L.; Cowburn, R. F.; Li, L.; Braak, H.; Alafuzoff, I.; Iqbal, K.; Iqbal, I. G.; Winblad, B.; Pei, J. J. Up-regulation of phosphorylated/activated p70 S6 kinase and its relationship to neurofibrillary pathology in Alzheimer’s disease. The American journal of pathology 2003, 163 (2), 591–607. DOI: 10.1016/S0002-9440(10)63687-5. Li, X.; Alafuzoff, I.; Soininen, H.; Winblad, B.; Pei, J. J. Levels of mTOR and its downstream targets 4E-BP1, eEF2, and eEF2 kinase in relationships with tau in Alzheimer’s disease brain. The FEBS journal 2005, 272 (16), 4211–4220. DOI: 10.1111/j.1742-4658.2005.04833.x. Tang, Z.; Bereczki, E.; Zhang, H.; Wang, S.; Li, C.; Ji, X.; Branca, R. M.; Lehtio, J.; Guan, Z.; Filipcik, P.; et al. Mammalian target of rapamycin (mTor) mediates tau protein dyshomeostasis: implication for Alzheimer disease. The Journal of biological chemistry 2013, 288 (22), 15556–15570. DOI: 10.1074/jbc.M112.435123.

(42) Wang, S.; Zhou, S. L.; Min, F. Y.; Ma, J. J.; Shi, X. J.; Bereczki, E.; Wu, J. mTOR-mediated hyperphosphorylation of tau in the hippocampus is involved in cognitive deficits in streptozotocin-induced diabetic mice. Metab Brain Dis 2014, 29 (3), 729–736. DOI: 10.1007/s11011-014-9528-1 From NLM.

(43) Sim, A. Y.; Choi, D. H.; Kim, J. Y.; Kim, E. R.; Goh, A. R.; Lee, Y. H.; Lee, J. E. SGLT2 and DPP4 inhibitors improve Alzheimer’s disease-like pathology and cognitive function through distinct mechanisms in a T2D-AD mouse model. Biomed Pharmacother 2023, 168, 115755. DOI: 10.1016/j.biopha.2023.115755 From NLM. Singh, A.; Tiwari, S.; Singh, S. Pirh2 modulates the mitochondrial function and cytochrome c-mediated neuronal death during Alzheimer’s disease. Cell Death Dis 2024, 15 (5), 331. DOI: 10.1038/s41419-024-06662-1 From NLM. Liu, S.; Fan, M.; Xu, J. X.; Yang, L. J.; Qi, C. C.; Xia, Q. R.; Ge, J. F. Exosomes derived from bone-marrow mesenchymal stem cells alleviate cognitive decline in AD-like mice by improving BDNF-related neuropathology. J Neuroinflammation 2022, 19 (1), 35. DOI: 10.1186/s12974-022-02393-2 From NLM.

(44) Yang, L.; Wang, D.; Zhang, Z.; Jiang, Y.; Liu, Y. Isoliquiritigenin alleviates diabetic symptoms via activating AMPK and inhibiting mTORC1 signaling in diet-induced diabetic mice. Phytomedicine 2022, 98, 153950. DOI: 10.1016/j.phymed.2022.153950 From NLM.

(45) Yao, D.; Bao, L.; Wang, S.; Tan, M.; Xu, Y.; Wu, T.; Zhang, Z.; Gong, K. Isoliquiritigenin alleviates myocardial ischemia-reperfusion injury by regulating the Nrf2/HO-1/SLC7a11/GPX4 axis in mice. Free Radic Biol Med 2024, 221, 1–12. DOI: 10.1016/j.freeradbiomed.2024.05.012 From NLM. Zeng, J.; Chen, Y.; Ding, R.; Feng, L.; Fu, Z.; Yang, S.; Deng, X.; Xie, Z.; Zheng, S. Isoliquiritigenin alleviates early brain injury after experimental intracerebral hemorrhage via suppressing ROS- and/or NF-κB-mediated NLRP3 inflammasome activation by promoting Nrf2 antioxidant pathway. J Neuroinflammation 2017, 14 (1), 119. DOI: 10.1186/s12974-017-0895-5 From NLM. Gao, Y.; Lv, X.; Yang, H.; Peng, L.; Ci, X. Isoliquiritigenin exerts antioxidative and anti-inflammatory effects via activating the KEAP-1/Nrf2 pathway and inhibiting the NF-κB and NLRP3 pathways in carrageenan-induced pleurisy. Food Funct 2020, 11 (3), 2522–2534. DOI: 10.1039/c9fo01984g From NLM. Qiu, M.; Ma, K.; Zhang, J.; Zhao, Z.; Wang, S.; Wang, Q.; Xu, H. Isoliquiritigenin as a modulator of the Nrf2 signaling pathway: potential therapeutic implications. Front Pharmacol 2024, 15, 1395735. DOI: 10.3389/fphar.2024.1395735 From NLM.

(46) Beurel, E.; Grieco, S. F.; Jope, R. S. Glycogen synthase kinase-3 (GSK3): regulation, actions, and diseases. Pharmacol Ther 2015, 148, 114–131. DOI: 10.1016/j.pharmthera.2014.11.016 From NLM.

(47) Pei, J.-J.; Braak, E.; Braak, H.; Grundke-Iqbal, I.; Iqbal, K.; Winblad, B.; Cowburn, R. F. Distribution of Active Glycogen Synthase Kinase 3β (GSK-3β) in Brains Staged for Alzheimer Disease Neurofibrillary Changes. Journal of Neuropathology & Experimental Neurology 1999, 58 (9), 1010–1019. DOI: 10.1097/00005072-199909000-00011 (acccessed 11/19/2024).

(48) Liu, L.; Li, Y.; Chen, G.; Chen, Q. Crosstalk between mitochondrial biogenesis and mitophagy to maintain mitochondrial homeostasis. Journal of Biomedical Science 2023, 30 (1), 86. DOI: 10.1186/s12929-023-00975-7.

(49) Norambuena, A.; Sagar, V. K.; Wang, Z.; Raut, P.; Feng, Z.; Wallrabe, H.; Pardo, E.; Kim, T.; Alam, S. R.; Hu, S.;, et al. Disrupted mitochondrial response to nutrients is a presymptomatic event in the cortex of the APP(SAA) knock-in mouse model of Alzheimer’s disease. Alzheimers Dement 2024. DOI: 10.1002/alz.14144 From NLM.

(50) Song, Z.; Zhang, Y.; Zhang, H.; Rajendran, R. S.; Wang, R.; Hsiao, C. D.; Li, J.; Xia, Q.; Liu, K. Isoliquiritigenin triggers developmental toxicity and oxidative stress-mediated apoptosis in zebrafish embryos/larvae via Nrf2-HO1/JNK-ERK/mitochondrion pathway. Chemosphere 2020, 246, 125727. DOI: 10.1016/j.chemosphere.2019.125727 From NLM.

(51) Lee, D. G.; Nam, B. R.; Huh, J. W.; Lee, D. S. Isoliquiritigenin Reduces LPS-Induced Inflammation by Preventing Mitochondrial Fission in BV-2 Microglial Cells. Inflammation 2021, 44 (2), 714–724. DOI: 10.1007/s10753-020-01370-2 From NLM.

(52) Wang, Z.; Jia, S.; Kang, X.; Chen, S.; Zhang, L.; Tian, Z.; Liang, X.; Meng, C. Isoliquiritigenin alleviates neuropathic pain by reducing microglia inflammation through inhibition of the ERK signaling pathway and decreasing CEBPB transcription expression. International Immunopharmacology 2024, 143, 113536. DOI: 10.1016/j.intimp.2024.113536.

(53) Nunomura, A.; Perry, G.; Pappolla, M. A.; Wade, R.; Hirai, K.; Chiba, S.; Smith, M. A. RNA oxidation is a prominent feature of vulnerable neurons in Alzheimer’s disease. J Neurosci 1999, 19 (6), 1959–1964. DOI: 10.1523/jneurosci.19-06-01959.1999 From NLM.

(54) Hwang, C. K.; Chun, H. S. Isoliquiritigenin isolated from licorice Glycyrrhiza uralensis prevents 6-hydroxydopamine-induced apoptosis in dopaminergic neurons. Biosci Biotechnol Biochem 2012, 76 (3), 536–543. DOI: 10.1271/bbb.110842 From NLM.

(55) Naia, L.; Shimozawa, M.; Bereczki, E.; Li, X.; Liu, J.; Jiang, R.; Giraud, R.; Leal, N. S.; Pinho, C. M.; Berger, E.;, et al. Mitochondrial hypermetabolism precedes impaired autophagy and synaptic disorganization in App knock-in Alzheimer mouse models. Mol Psychiatry 2023, 28 (9), 3966–3981. DOI: 10.1038/s41380-023-02289-4 From NLM.

(56) Terada, T.; Obi, T.; Bunai, T.; Matsudaira, T.; Yoshikawa, E.; Ando, I.; Futatsubashi, M.; Tsukada, H.; Ouchi, Y. In vivo mitochondrial and glycolytic impairments in patients with Alzheimer disease. Neurology 2020, 94 (15), e1592–e1604. DOI: 10.1212/wnl.0000000000009249 From NLM.

(57) Ruegsegger, G. N.; Manjunatha, S.; Summer, P.; Gopala, S.; Zabeilski, P.; Dasari, S.; Vanderboom, P. M.; Lanza, I. R.; Klaus, K. A.; Nair, K. S. Insulin deficiency and intranasal insulin alter brain mitochondrial function: a potential factor for dementia in diabetes. Faseb j 2019, 33 (3), 4458–4472. DOI: 10.1096/fj.201802043R From NLM.

(58) da Silva, E. M. G.; Fischer, J. S. G.; Souza, I. L. S.; Andrade, A. C. C.; Souza, L. C. E.; Andrade, M. K.; Carvalho, P. C.; Souza, R. L. R.; Vital, M.; Passetti, F. Proteomic Analysis of a Rat Streptozotocin Model Shows Dysregulated Biological Pathways Implicated in Alzheimer’s Disease. Int J Mol Sci 2024, 25 (5). DOI: 10.3390/ijms25052772 From NLM.

(59) Denzer, I.; Münch, G.; Pischetsrieder, M.; Friedland, K. S-allyl-l-cysteine and isoliquiritigenin improve mitochondrial function in cellular models of oxidative and nitrosative stress. Food Chemistry 2016, 194, 843–848. DOI: 10.1016/j.foodchem.2015.08.052.

(60) Huang, Y.; Sun, J.; Li, S.; Shi, Y.; Yu, L.; Wu, A.; Wang, X. Isoliquiritigenin mitigates intervertebral disc degeneration induced by oxidative stress and mitochondrial impairment through a PPARγ-dependent pathway. Free Radic Biol Med 2024, 225, 98–111. DOI: 10.1016/j.freeradbiomed.2024.10.001 From NLM.

(61) Zappa Villar, M. F.; López Hanotte, J.; Falomir Lockhart, E.; Trípodi, L. S.; Morel, G. R.; Reggiani, P. C. Intracerebroventricular streptozotocin induces impaired Barnes maze spatial memory and reduces astrocyte branching in the CA1 and CA3 hippocampal regions. J Neural Transm (Vienna) 2018, 125 (12), 1787–1803. DOI: 10.1007/s00702-018-1928-7 From NLM.

(62) Mehla, J.; Pahuja, M.; Gupta, Y. K. Streptozotocin-Induced Sporadic Alzheimer’s Disease: Selection of Appropriate Dose. Journal of Alzheimer’s Disease 2012, 33 (1), 17–21. DOI: 10.3233/JAD-2012-120958 (acccessed 2024/10/08). Canet, G.; Gratuze, M.; Zussy, C.; Bouali, M. L.; Diaz, S. D.; Rocaboy, E.; Laliberté, F.; El Khoury, N. B.; Tremblay, C.; Morin, F.; et al. Age-dependent impact of streptozotocin on metabolic endpoints and Alzheimer’s disease pathologies in 3xTg-AD mice. Neurobiol Dis 2024, 198, 106526. DOI: 10.1016/j.nbd.2024.106526 From NLM.

(63) Kong, Y.; Cao, L.; Xie, F.; Wang, X.; Zuo, C.; Shi, K.; Rominger, A.; Huang, Q.; Xiao, J.; Jiang, D.;, et al. Reduced SV2A and GABAA receptor levels in the brains of type 2 diabetic rats revealed by [18F]SDM-8 and [18F]flumazenil PET. Biomedicine & Pharmacotherapy 2024, 172, 116252. DOI: 10.1016/j.biopha.2024.116252.

