## Supplemental file for "Isoliquiritigenin attenuated cognitive impairment, cerebral tau phosphorylation and oxidative stress in a streptozotocin-induced mouse model of Alzheimer’s disease"

**Supplementary Table 1 Antibody and chemical used in the study**

| **Antibody** | **Host** | **WB/IF Dilution** | **Sources** | **Catalog No** |
| --- | --- | --- | --- | --- |
| NeuN | r | 1:400 | Abcam | #ab177487 |
| Anti-DNA/RNA Damage antibody [15A3] | m | 1:50 | Abcam | #ab62623 |
| p-tau396 | r | 1:1000 | Thermo Fisher | #35-5300 |
| Tau5 | r | 1:1000 | Abcam | #ab80579 |
| Mfn1 | m | 1:500 | Santa Cruz | #sc-166644 |
| Mfn2 | m | 1:500 | Santa Cruz | #sc-100560 |
| DRP1 | r | 1:1000 | Abcam | #ab184247 |
| p-mTOR | m | 1:1000 | Cell Signaling Technology | #5536s |
| t-mTOR | r | 1:1000 | Cell Signaling Technology | #2983s |
| p-GSK-3β (Ser9) | r | 1:1000 | Cell Signaling Technology | #9323s |
| t-GSK-3β | r | 1:1000 | Cell Signaling Technology | #12456s |
| p-Erk1/2 (T202/Y204) | r | 1:1000 | Cell Signaling Technology | #4370S |
| t-Erk1/2 | r | 1:1000 | Cell Signaling Technology | #4695S |
| PSD95 | r | 1:1000 | Abcam | #ab238135 |
| SNAP25 | r | 1:2000 | Abcam | #ab108990 |
| β-tubulin | r | 1:10000 | Abcam | #ab179511 |
| β-actin | r | 1:20000 | Genetex | #GTX109639 |
| **Chemical** |  |  | **Sources** | **Catalog No** |
| Isoliquiritigenin |  |  | Medchem express | HY-N0102 |
| Trypan blue |  |  | Beyotime | ST2780-5g |
| Streptozotocin |  |  | Sigma Aldrich | 18883-66-4 |

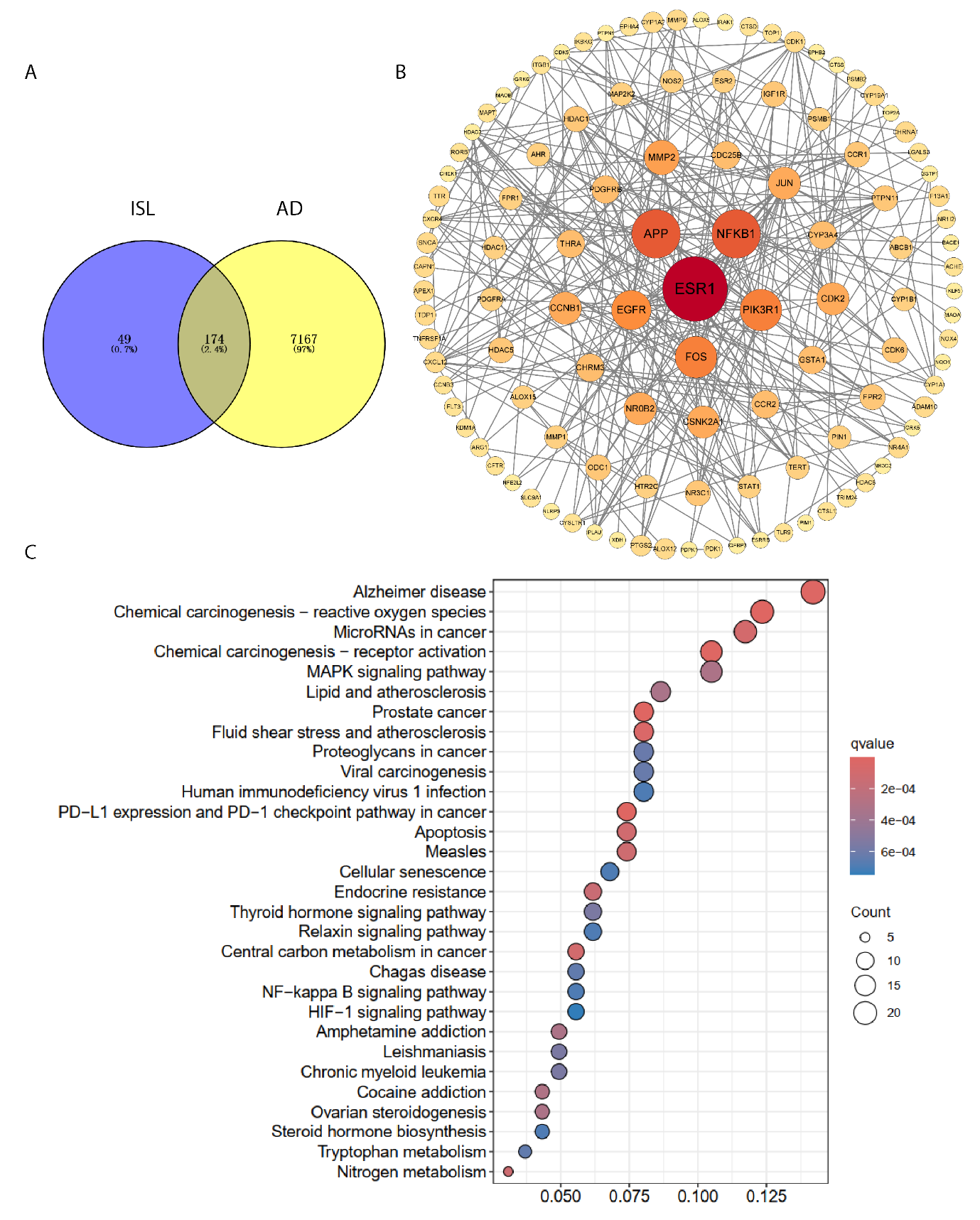

**SFig. 1 isoliquiritigenin target the activation of Alzheimer’s disease, reactive oxygen species, the MAPK pathway based on network pharmacology.** **(A)** Drug-disease related targets. (**B**) Core targets and network interactions. Network pharmacology results of action of isoliquiritigenin in AD. (**C**) KEGG enrichment analysis of treatment-related AD targets.
